# Strategies of host resistance to pathogens in spatially structured populations: An agent-based evaluation

**DOI:** 10.1101/423822

**Authors:** Christophe Boëte, Morgan Seston, Mathieu Legros

## Abstract

There is growing theoretical evidence that spatial structure can affect the ecological and evolutionary outcomes of host-parasite interactions. Locally restricted interactions have been shown in particular to affect host resistance and tolerance. In this study we investigate the evolution of several types of host disease resistance strategies, alone or in combination, in spatially structured populations. We construct a spatially explicit, individual-based stochastic model where hosts and parasites interact with each other in a spatial lattice, and interactions are restricted to a given neighbourhood of varying size. We investigate several host resistance strategies, including constitutive (expressed in all resistant hosts), induced (expressed only upon infection), and combinations thereof. We show that the specific resistance mechanism against a pathogen as well as the size of the neighbourhood both affect resistance spread and parasite dynamics, an effect modulated by the level of the cost of resistance. Our results shed new light on the dynamics of disease resistance in a spatially-structured host-pathogen system, and illustrate the conditions in which a variety of resistance mechanisms can be maintained, including when they are costly. Overall these results advance our theoretical understanding of the evolutionary dynamics of disease resistance, a necessary step to elaborate more efficient and sustainable strategies for disease management.

## 1. Background

Host resistance to pathogens presents an obvious evolutionary advantage and, because of the ubiquity of parasitism in the living world, such traits are typically and pervasively under strong, positive selection. Resistance, however, is usually associated with a fitness cost for the host, and therefore also subject to negative selective pressure. Such costs of resistance have been demonstrated in many organisms ranging from bacteria [1] to insects [2, 3] and plants [4]. The balance between these selective forces drives the evolution of resistance in host-parasite systems.

There is now growing evidence that the spatial structure of populations can affect this selective balance, and therefore affect the ecological outcomes of host-parasite interactions, and ultimately the epidemiological dynamics of diseases. In theoretical models [5], locally restricted interactions between hosts and pathogens have been shown to favour host resistance when compared to well-mixed population models [6]. In addition, the evolution of host tolerance has been shown to be restricted in spatially explicit population models [7].

All the aforementioned factors (pathogen pressure, costs of resistance, spatial population structure) paint the picture of a complex selective landscape for host resistance to disease. Consequently, depending on the specific host and pathogen species of interest, as well as ecological and epidemiological characteristics of the considered system, it is likely that the actual selection pressures would differ substantially and, as a corollary, that different resistance strategies would be favoured by selection. Generally speaking, resistance strategies can differ in the mechanism itself (e.g. specific immune pathways), in the quantitative investment in the response, or in the timing of the response (e.g. constitutive expression of immune components versus induced expression in response to an infection).

The purpose of this study is therefore to further characterize the dynamics of the evolution of resistance strategies in a spatially-structured host-parasite system, with a specific focus on the different types of resistance (and combinations thereof) that a host can possibly mount and the corresponding impacts of the associated resistance costs.

To that end, we present in this article a spatially explicit, individual-based stochastic model, simulating host and pathogen populations and their interactions. The spatial architecture of this model allows us to restrict to a given neighbourhood (of varied size) the biological and ecological processes in the model, such as pathogen transmission dynamics, host reproduction, and migration, and therefore to control the level of spatial structure in the population. Within this (more or less) spatially restricted framework we examine the fate of a variety of host resistance strategies against a generic pathogen. In particular we consider whether resistance (and its associated costs) is constitutively expressed in resistant hosts or induced by the response to an infection (and therefore not expressed in uninfected hosts). We also distinguish between responses aimed at stopping forward transmission versus clearing an existing pathogen in an infected host.

In the following sections, we first present the details of the model features and parameterization. We then present the result of model simulations that examine resistance strategies based on constitutive response, induced responses affecting transmission or recovery, or a mix thereof, all under various levels of spatial population structure (i.e. locally restricted interactions). We demonstrate that the evolution of resistance in a spatial setting depends on the type of resistance and the conditions in which it is mounted by the host, along with the costs associated with each type of resistance. We conclude by discussing the evolutionary consequences of these observations for host-pathogen and disease systems.

## 2. Model

### 2.1 General model characteristics

We introduce in this study a probabilistic cellular automaton, a stochastic, spatially-explicit individual-based model. The population is described as a square, homogeneous 100×100 lattice of individual sites, each of which may be empty or occupied by a single (infected or uninfected) host. To eliminate edge effects, we assume that the lattice has a toric topology, i.e. sites on a given edge are connected to sites on the opposite edge. The source code for the simulation is available on request.

### 2.2. Host genotypes and resistance

Hosts can be classified into two main genotypes according to their ability to respond to a pathogen. Susceptible hosts are completely vulnerable to pathogens and not able to mount any response to infection. Resistant hosts are able, through immune responses, to resist infection to some extent (see details below, depending on the mode of resistance and the investment in the response).

We consider a variety of response strategies that resistant hosts can mount against pathogens. First, we distinguish between constitutive and inducible immune responses. A constitutive response is expressed by all resistant hosts regardless of their infection status, whereas an inducible response is only expressed upon infection. A given host can express a mixture of these two strategies, where the ratio of investment in constitutive response is defined as r1 (in other words, if r1=1, then the response is strictly constitutive, and conversely, strictly inducible when r1= 0).

The constitutive immune response, being expressed at all times, is considered here equivalent to a barrier to infection. Thus, in our model, the constitutive immune response affects the probability, for a given uninfected host, of becoming infected. An inducible response, on the other hand, being only expressed upon infection, can only affect the pathogen dynamics within an infected host, or its ability to infect other hosts. Consequently, we distinguish in our model two modes of action for an inducible response: one affecting the clearance rate of the infection, the other affecting the forward transmissibility of the parasite to other hosts. The investment in one or the other is determined by a factor called r2. If r2= 1, the response affects only the clearance, whereas only transmissibility is reduced when r2=0. This means that inducible strategies can exist along a continuum going from purely inward strategies targeted at the infected individual itself (i.e. focused on clearing the infection, r2=1) to outward strategies targeted at its interactions with neighbours (i.e. focused on blocking forward transmission, r2=0, where the resistant individual gets no direct benefit).

Resistance is also characterized by a given efficacy (*eff1* or *eff2* for constitutive and inducible responses respectively) negatively affecting the parasite fitness in a manner depending of the type of resistance. When considering one single response, the efficacy of resistance is always total (equal to 1) whereas in the case of multiple responses the efficacy is directly related to the ratio of the investment in the given responses.

### 2.3. Model dynamics

At each time step, the model is iterated through the following processes in the following order (each detailed below): reproduction, movement, death and change in infectious status (infection, recovery). In this agent-based framework, local interactions are restricted to individuals within a given neighbourhood. In this study we use van Neumann neighbourhoods, with sizes (*sz*) varying between 1 and 5, where the size of the neighbourhood corresponds to the maximum Manhattan distance (distance between two points in a grid based on a strictly horizontal and/or vertical path) between neighbours. The higher the value is, the less viscous is the population, i.e. closer to a non-structured, well-mixed system. This type of spatial structuration and the associated levels of population viscosity have indeed been shown to influence the evolution of a variety of life-history traits in host parasite systems. Pathogens in spatially structured population tend to evolve lower transmission rate and virulence as shown in theoretical approaches (Boots and Sasaki, 1999; Kamo et al. 2007; Lion and Boots, 2010) or via empirical studies (Kerr et al. 2006; Boots and Mealor 2007), whereas localized transmission and host reproduction can select for a higher host resistance (Best et al. 2011).

#### 2.3.1. Reproduction

Each individual can give birth by asexual reproduction to one uninfected (no vertical transmission) offspring per time step. This offspring is of the same genotype and expresses the same type of immune response (if any). The nominal birth rate, denoted *rH*, is the same for infected and uninfected individuals as we assume that fecundity is not affected by the infection (and that pathogen virulence only affects survival, see below).

In a similar manner as done in previous studies [6, 8] resistance is assumed to be associated with a cost leading to a reduced fecundity of the resistant individuals by a factor (1-cR), where cR is the cost of resistance.

All resistant individuals pay the cost of a constitutive resistance, and the reduction of fecundity is defined as:

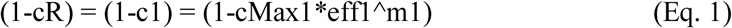

Concerning the induced resistance, only the infected resistant individuals (as resistance is only expressed during infection) pay the associated cost. The reduction of fecundity is then defined as:

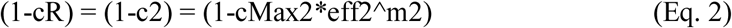

When both responses are combined then the cost paid by the infected individuals is assumed to be multiplicative:

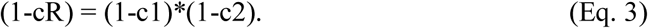

In Eq. 3 and 4, both m1 and m2 represent the departure from linearity. The higher their value is, the lower the cost is at low levels of investment, i.e. efficacy. A list of model parameters, along with their definition and the range of explored values, is given in Table 1. Table 2 recapitulates the host life history traits for various types of hosts based on these parameters.

**Table 1:** Definitions and range of values of the model parameters.

| Parameter | Definition | Range of values |
| --- | --- | --- |
| cMax1 | Maximal cost of resistance of the constitutive immune response | 0 - 1 |
| cMax2 | Maximal cost of resistance of the inducible immune response | 0 - 1 |
| r1 | Ratio of the investment between the constitutive immune response and the inducible one(s) | 0 - 1 |
| r2 | Ratio of the investment between the inducible recovery-boosting and inducible transmission-blocking responses | 0 - 1 |
| rH | Birth rate of a susceptible (S) individual | 0.02 - 0.08 |
| $\beta_0$ | Transmission rate from an infected susceptible (I) individual | 0.125 - 0.5 |
| $\alpha$ | Virulence of the parasite (additional mortality) | 0.125 - 0.5 |
| $\mu$ | Death rate of a susceptible (S) individual | 0.1 |
| $\gamma$ | Recovery rate of the infected susceptible (I) individual | 0.1 |
| eff1 | Efficacy of the constitutive resistance | 0 - 1 |
| eff2 | Efficacy of the inducible resistance | 0 - 1 |
| sz | Neighbourhood size | 1 - 5 |
| m1 - m2 | Departure from linearity | 1.4 |

**Table 2:** Life-history traits of the different types of hosts and epidemiological parameters of the model

|  | Susceptible<br>Uninfected | Susceptible<br>Infected | Resistant Uninfected | Resistant Infected |
| --- | --- | --- | --- | --- |
| Reproduction | $rH$ | $rH$ | $rH(1-c1)$ | $rH*(1-c1)*(1-c2)$ |
| Mortality | $\mu$ | $\mu(1+\alpha M)$ | $\mu$ | $\mu*(1+\alpha)$ |
| Transmission Rate | 0 | $\beta_0$ | 0 | $\beta_0*(1-eff2*(1-r2))$ |
| Clearance | - | $\gamma$ | - | $1-(1-\gamma)*(1-eff2*r2)$ |
| Cost of the Constitutive Resistance (c1) | - | - | $cMax1*eff1^{m1}$ | $cMax1*eff1^{m1}$ |
| Cost of the Inducible Resistance (c2) | - | - | - | $cMax2*eff2^{m2}$ |

Host reproduction is implemented as a two-step process. First, there is a random choice of a number between 0 and 1 (uniform distribution). If its value is lower than the reproduction rate, then there is a second draw choosing a random site (with equiprobability for each site) in the neighbourhood. If it is an empty one, then the offspring occupies this site, otherwise there is no reproduction for the host at this given round. The reproduction process is therefore density-dependent, and the nominal birth rate rH represents the optimal reproduction rate for an isolated individual. During this process, it is obvious that the density-dependence advantages the first hosts being selected by the algorithm. To avoid systematically favouring specific locations in the lattice, hosts are treated in a random order at each time step.

#### 2.3.2. Movement

A Bernoulli trial determines the occurrence of movement for a given host with a probability given by the movement rate. If movement occurs, a destination site is chosen at random in the neighbourhood. If it is an empty site, then the host moves, otherwise the host stays in the site. When movement occurs, the genotype of the host and its infection status remain unchanged.

#### 2.3.3. Death

Individuals are removed from the population at a mortality rate that depends on the infectious status. Each host may die at each time step based on the outcome of a Bernoulli trial with a probability given by the mortality rate. At the end of this step, a snapshot of the grid is done in order to record all information needed to evaluate the next processes.

#### 2.3.4. State change: Infection

In the spatial model, the parasite can be transmitted from an infected host to uninfected hosts among its neighbours in a frequency-dependent manner. For a given uninfected host, the rate of infection depends on the presence of infected hosts in the neighbourhood. Each infected neighbour’s ability to transmit the parasite is characterized by a probability of infection, denoted *β*, that is a measure of the parasite’ success in transmission. If this infected host has a resistant genotype with an inducible response acting on transmission (r2<1), the probability of infection from this particular host is reduced by a factor (1-eff2^∗^(1-r2)).

For a given uninfected host, the probability of infection is then calculated as the probability of successful transmission by at least one of its infected neighbours. A random number between 0 and 1 (uniform distribution) is then drawn and if it is lower than this probability of infection, transmission occurs. Resistant individuals expressing a constitutive response with efficacy eff1 have a probability to become infected that is reduced by a factor (1-eff1). Resistant individuals expressing an inducible response, on the other hand, have the same probability to be infected than the susceptible ones. Table 3 describes the pairwise probability of infection of a host X by a host Y.

**Table 3:**
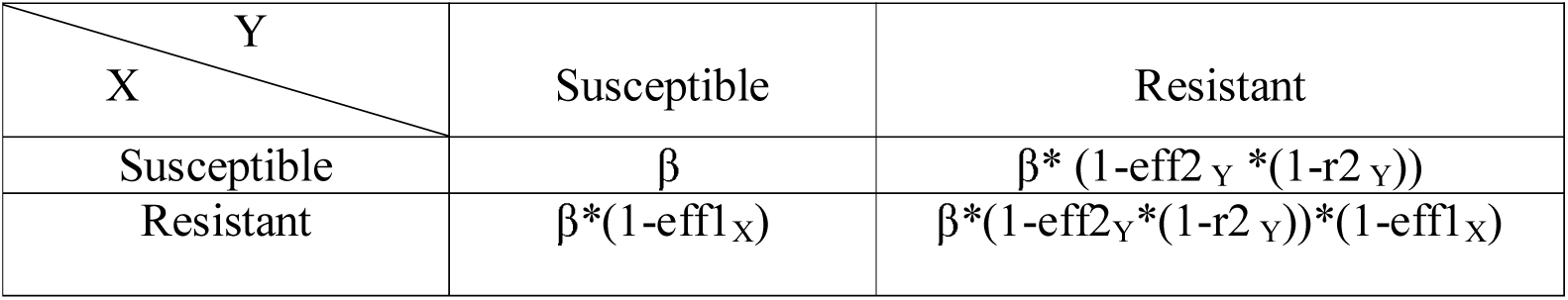
Probability of infection of an uninfected host X by an infected host Y.

#### 2.3.5. State change: Recovery

Individuals can recover from infection with a rate *γ*. The recovery rate can be affected by the investment in the inducible defence and in this latter case, the recovery rate becomes:

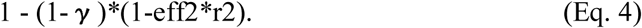

In a similar manner as for the previous events, if the recovery rate is lower than a randomly drawn number chosen between 0 and 1, then the individual remains infected.

#### 2.3.6. Simulations

In each simulation, the lattice is initially populated with 1000 individuals of the susceptible genotype including 20% of infected, all randomly distributed between sites. The simulation is first run for 1000 time steps to allow the resident population to equilibrate. 100 non-infected invading resistant hosts are then introduced into sites randomly selected among the available empty sites in the lattice. The simulation is run for 250 000 iterations or stopped earlier if one genotype reaches fixation. Except when stated otherwise, five replicates are done for each set of parameters.

## 3. Results

In the first two scenarios, we only explored the effect of space on the spread of one type of resistance alone, first the constitutive one, then the inducible one. To do so the r1 parameter was respectively set to 1 then 0. We varied the cost associated with each type of resistance, as well as the size of the neighbourhood, in order to measure their combined impact on the spread of resistance in the host population as well as on the epidemiology of the disease.

### 3.1. Constitutive resistance alone (r1=1)

As mentioned earlier constitutive resistance is associated with a fixed cost (paid by both infected and non-infected individuals). As there is only a single type of resistance involved, the value of this cost is cMax1. We observe that for all neighbourhood sizes a moderately high cost (cost>0.1) precludes the spread of resistance altogether (Figure 1). Fixation of the resistant genotype can only occur when resistance is not costly. In those instances, the parasite is then always eliminated from the population at the end of the corresponding simulations. The parasite can also be eliminated from the population for a range of low, non-zero values of cost, in which case the resistant genotype is then unsurprisingly lost as a consequence of this cost.

**Figure 1A.**
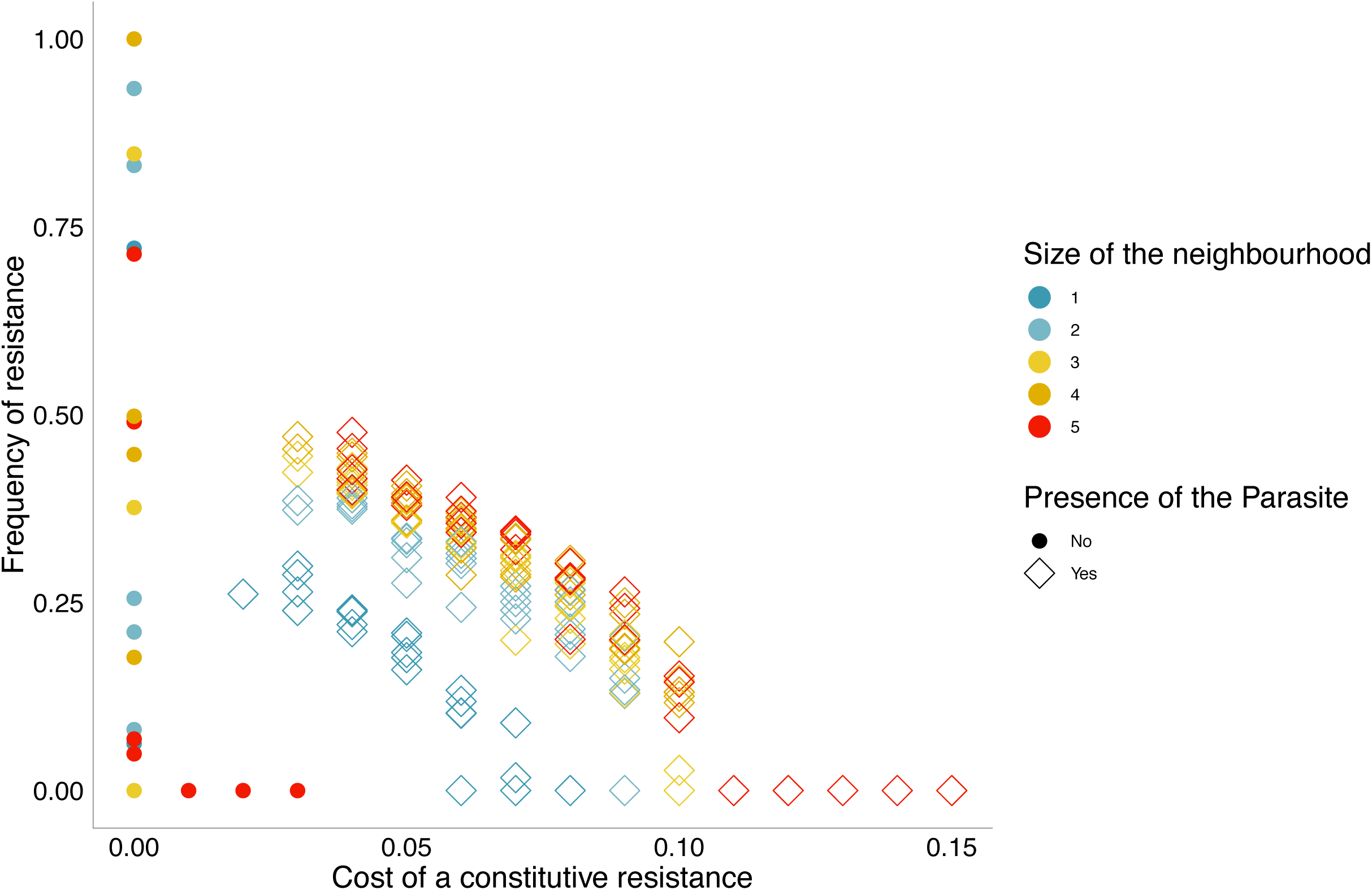
Effect of the fitness cost of resistance (cMax1) and of the size of neighbourhood on the spread of a constitutive resistance against a pathogen and on the parasite presence at equilibrium. The parameters are the following ones: transmission rate β=0.25, virulence of the pathogen alpha=0.25, mortality μ=0.01, recovery rate γ=0.1, reproduction rate rH=0.02, efficacy of resistance eff1=1, departure from linearity m1=1.4, level of investment in the constitutive resistance r1=1.

The size of the neighbourhood affects both resistance spread and parasite dynamics. We observe that, for a given cost, resistance can spread to higher frequencies when neighbourhood size is larger (Figure 1A). These results are confirmed with a simple statistical analysis (Table 4)

**Table 4.** Logistic analysis of the spread (presence/ absence) of a costly constitutive defence response in different spatial contexts.

| Source | d.f. | Log-likelihood $\chi^2$ | P-value |
| --- | --- | --- | --- |
| Size of the neighbourhood | 4 | 9.42 | 0.052 |
| Cost of resistance | 1 | 47.39 | <0.0001 |
| Cost of resistance x Size of the neighbourhood | 4 | 1.917 | 0.751 |
| Replicate | 4 | 0.864 | 0.929 |

Additionally, we observe that in larger neighbourhoods, parasites can be eliminated from the population at higher values of cost of resistance. It is noteworthy that parasites are eliminated at a higher speed for low values of the cost of resistance (Figure 1B).

**Figure 1B.**
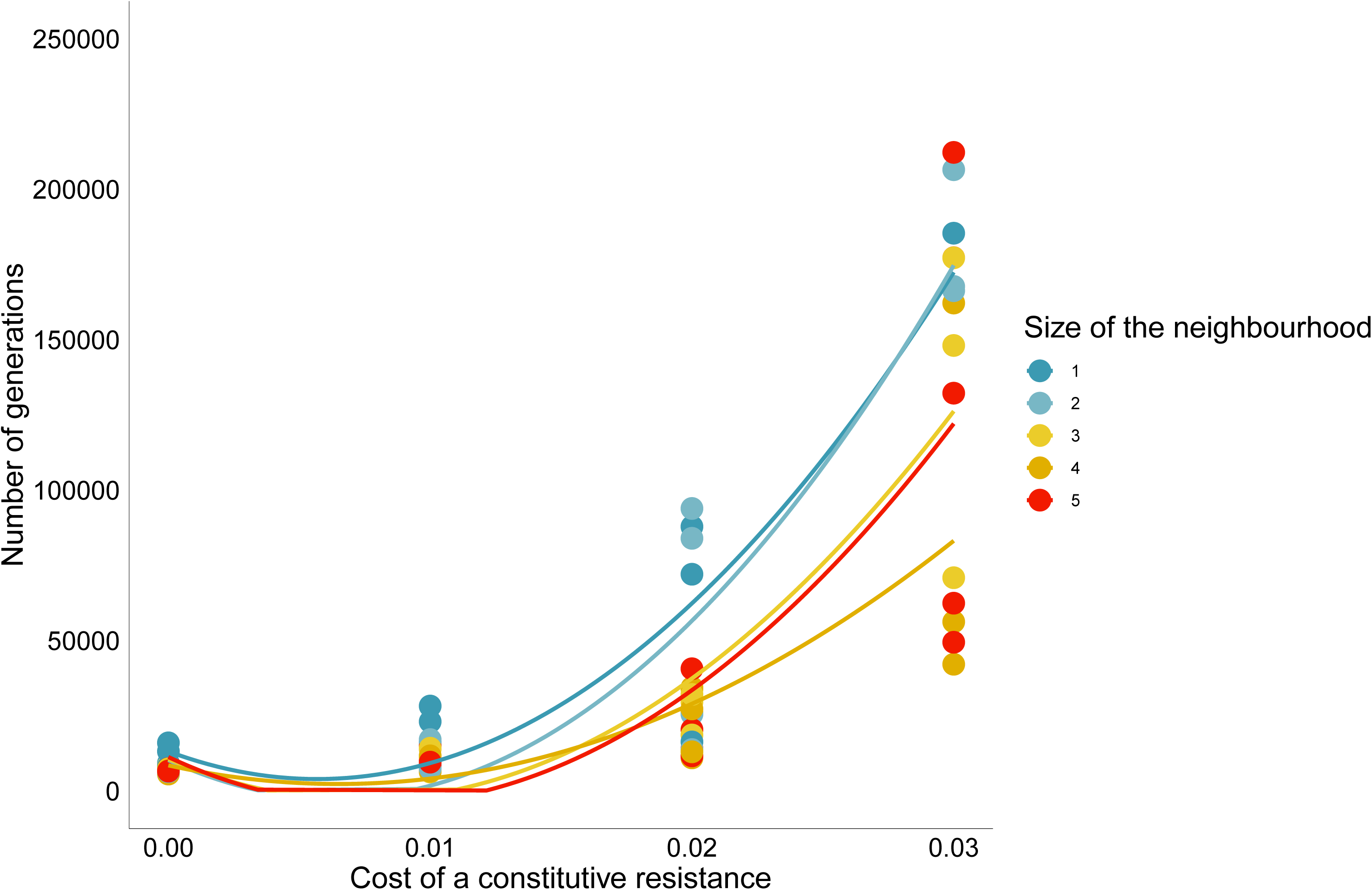
Effect of the fitness cost of a constitutive resistance against a pathogen and of the size of neighbourhood on the speed of elimination of a parasite. The parameters are the following ones: transmission rate β=0.25, virulence of the pathogen alpha=0.25, mortality μ=0.01, recovery rate γ=0.1, reproduction rate rH=0.02, efficacy of resistance eff1=1, departure from linearity m1=1.4, level of investment in the constitutive resistance r1=1. The regression lines in the graphs are built for each size of neighbourhood with the points for which the parasite is still present.

### 3.2. Inducible defences alone (r1=0)

As mentioned in the Methods, in the case of inducible defence, we explore two distinct mechanisms, either blocking forward transmission (reducing β) or boosting recovery (increasing γ). The ratio of the investment in each mechanism is given by r2, where r2=0 corresponds to an exclusive investment in transmission-blocking response, and r2=1 to an exclusive investment in recovery-boosting response. We first examine each of these responses separately.

#### 3.2.1. Transmission-blocking response (r1=0 and r2=0)

The size of the neighbourhood has a strong impact on the spread of this type of resistance (Table 5). We observe that when the cost of resistance increases, resistance spreads more easily in smaller neighbourhoods (S1 Figure).

**Table 5.** Logistic analysis of the spread (presence/ absence) of a costly inducible transmission-blocking defence response in different spatial contexts.

| Source | d.f. | Log-likelihood $\chi^2$ | P-value |
| --- | --- | --- | --- |
| Size of the neighbourhood | 4 | 38.92 | <0.0001 |
| Cost of resistance | 1 | 2.60 | 0.1071 |
| Cost of resistance x Size of the neighbourhood | 4 | 7.46 | 0.11 |
| Replicate | 4 | 2.73 | 0,60 |

The resistant genotype can be maintained in our simulations with relatively high values of the cost of resistance only in the smallest neighbourhood (size 1 and 2) and only in simulations where the parasites are eventually lost from the population. In the largest neighbourhoods resistance cannot become fixed and can only spread when it is not costly, while parasites can maintain especially when resistance is costly (Table 6).

**Table 6.** Logistic analysis of the presence of the parasite after the introduction of a costly inducible transmission-blocking defence response in different spatial contexts.

| Source | d.f. | Log-likelihood $\chi^2$ | P-value |
| --- | --- | --- | --- |
| Size of the neighbourhood | 4 | 50.32 | <0.0001 |
| Cost of resistance | 1 | 10.20 | 0.0014 |
| Cost of resistance x Size of the neighbourhood | 4 | 6.78 | 0.15 |
| Replicate | 4 | 4.33 | 0,36 |

#### 3.2.2. Recovery-boosting response (r1=0 and r2=1)

With this type of inducible defence, we observe that the interaction between the cost of resistance and the size of the neighbourhood significantly affects the spread of resistance. To a lesser extent, the size of the neighbourhood also affects the spread of resistance (Table 7).

**Table 7.** Logistic analysis of the spread (presence/ absence) of a costly inducible recovery-boosting response in different spatial contexts.

| Source | d.f. | Log-likelihood $\chi^2$ | P-value |
| --- | --- | --- | --- |
| Size of the neighbourhood | 4 | 8.48 | 0.075 |
| Cost of resistance | 1 | 0.66 | 0.4183 |
| Cost of resistance x Size of the neighbourhood | 4 | 90.47 | <0,0001 |
| Replicate | 4 | 6.59 | 0.16 |

Thus, when resistance is associated with a low cost, it is more likely to spread in populations with a low structuration (high sizes of neighbourhood). This trend is reversed in highly structured populations where resistance is then more likely to be maintained when the cost is high. Interestingly, this observed pattern differs from the one observed for the other type of inducible response, i.e. the transmission-blocking one (see § 3.2.1).

This is likely associated with the epidemiological dynamics in such populations, as we observe that parasites persist only for high levels of cost and even more in large neighbourhoods (Table 8) and only when resistance is eliminated. In addition, the lower frequency of resistant uninfected hosts at very high levels of cost and for large neighbourhoods is associated with the higher likelihood of maintenance of the parasite under those circumstances (S2 Figure).

**Table 8.** Logistic analysis of the presence of the parasite of a costly inducible recovery-boosting response in different spatial contexts.

| Source | d.f. | Log-likelihood $\chi^2$ | P-value |
| --- | --- | --- | --- |
| Size of the neighbourhood | 4 | 4.96 | 0.29 |
| Cost of resistance | 1 | 27.96 | <0,0001 |
| Cost of resistance x Size of the neighbourhood | 4 | 10.29 | 0.036 |
| Replicate | 1 | 0.87 | 0,93 |

#### 3.2.3. Mixing two inducible defence mechanisms: transmission-blocking and recovery-boosting (r1=0 and 0 < r2 < 1)

We now consider the case of a fluctuating investment of the resistant hosts between the two inducible responses, the transmission-blocking and the recovery-boosting one. In other words, this means that r2 can take values between 0 and 1.

With such mixed strategies, resistance can spread in the population and reach a high frequency for limited values of the cost especially for neighbourhoods of limited size (Figure 2)

**Figure 2.**
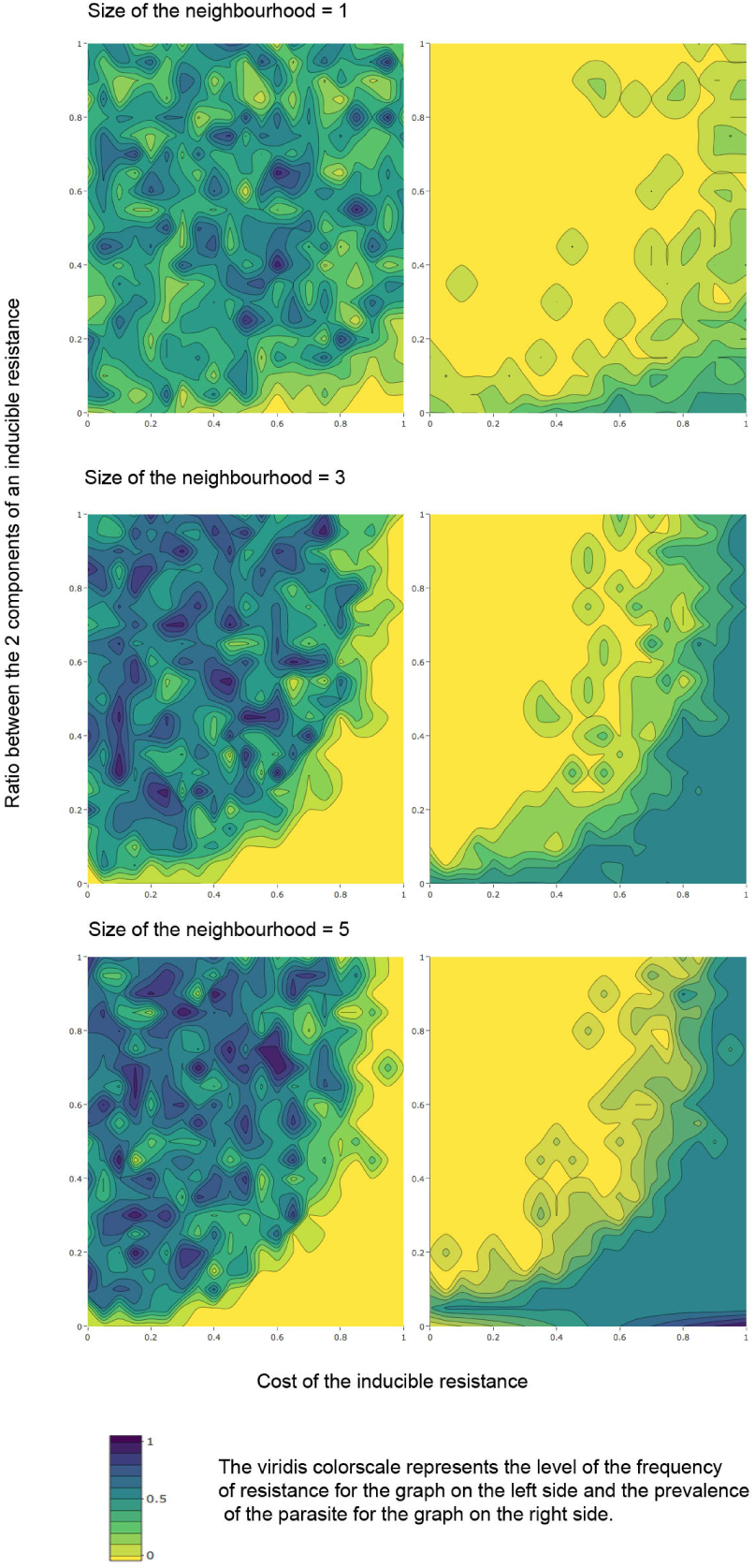
Effect the cost and the ratio between 2 components of an inducible resistance having a transmission-blocking and a recovery effect against a pathogen on its spread in different spatial structure and on the parasite prevalence at equilibrium. The parameters are the following ones: transmission rate β=0.25, virulence of the pathogen alpha=0.25, mortality μ=0.01, recovery rate γ=0.1, reproduction rate rH=0.02, efficacy of resistance eff1=1, departure from linearity m1=1.4. The level of investment in the 2 components of the response (r2) varies between 0 and 1. (Note that this is set of simulations was established with 4 replicates and for 3 levels of neighbourhood).

Given that the two considered inducible defence mechanisms exhibited contrasting patterns of sensitivity to levels of spatial population structure, it is especially interesting to observe that a strategy mixing these two mechanisms generally favours costly resistance in more highly structured populations (i.e. smaller neighbourhoods). Once the cost of resistance increases, it appears that a ratio biased towards the recovery response is favoured over the transmission-blocking response.

For a small neighbourhood, the cost, the ratio and their interaction are affecting the presence of resistance and an intermediate strategy is the one reaching the highest frequency of resistant hosts when the cost increases (Table 9; Figure 3). For larger neighbourhoods, only the cost and the ratio alone influence the presence of resistance (Table 9) and the selected strategy is the one favouring recovery response (Figure 3).

**Figure 3.**
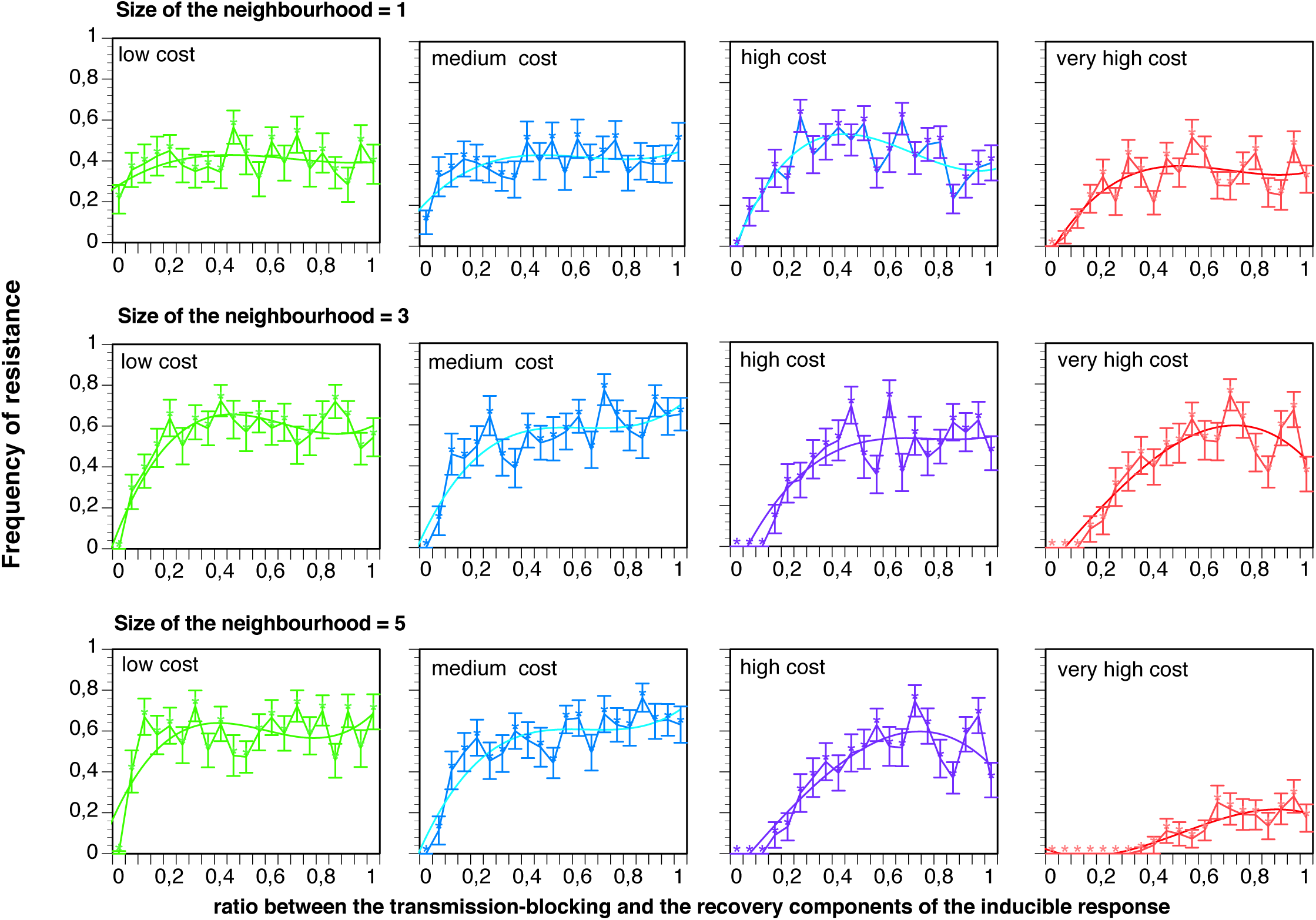
Effect the cost of an inducible resistance with a transmission-blocking and a recovery component against a pathogen on its spread in different spatial structure. The parameters are the following ones: transmission rate β=0.25, virulence of the pathogen alpha=0.25, mortality μ=0.01, recovery rate γ=0.1, reproduction rate rH=0.02, efficacy of resistance eff1=1, departure from linearity m1=1.4. The level of investment in the 2 components of the response (r2) varies between 0 and 1. The categories of the cost are the following ones: low [0-0.25[, medium [0.25-0.5[; high [0.5-0.75[ and very high [0.75-1]. (Note that this is set of simulations was established with 4 replicates and for 3 levels of neighbourhood).

**Table 9:** Logistic analysis of the spread (presence/ absence) of a costly inducible response mixing a recovery boosting and a transmission-blocking component in different spatial contexts.

| Source | d.f. | Log-likelihood $\chi^2$ | P-value |
| --- | --- | --- | --- |
| Size of the neighbourhood | 2 | 17.75 | 0.0001 |
| Ratio between the recovery-boosting and a transmission-blocking components (r2) | 1 | 45.0 | <0.0001 |
| Cost of resistance | 1 | 14.54 | 0.0001 |
| r2 x Cost of resistance | 1 | 17.89 | <0.0001 |
| r2 x Size of the neighbourhood | 2 | 94.23 | <0.0001 |
| Cost of resistance x Size of the neighbourhood | 2 | 313.24 | <0.0001 |
| Cost of resistance x Size of the neighbourhood x r2 | 2 | 19.69 | <0.0001 |
| replicate | 3 | 1.37 | 0,71 |

### 3.3. Mixing constitutive and inducible defence strategies (0 < r1 < 1)

We now consider the presence of resistant individuals able to invest in both the constitutive resistance and one of the inducible response, in other words scenarios where 0 < r1 <1.

#### 3.3.1. Mixing a constitutive and a recovery-boosting response (0 < r1 < 1 and r2=1)

In this case the host can invest in both a constitutive defence and an inducible response increasing recovery (r2=1). Interestingly the highest values of the frequency of resistance are obtained for an intermediate investment in both responses. While the ratio alone is not significantly affecting the presence of resistance, the size of the neighbourhood and its interaction with the ratio do have a strong impact (Table 9).

**Table 9.** Logistic analysis of the spread (presence/ absence) of a response with a constitutive component and a recovery-boosting one in different spatial contexts.

| Source | d.f. | Log-likelihood $\chi^2$ | P-value |
| --- | --- | --- | --- |
| Size of the neighbourhood | 4 | 11.66 | 0.02 |
| Ratio between the constitutive and the inducible recovery boosting response ($r_1$) | 1 | 1.25 | 0.27 |
| $r_1 \times$ Size of the neighbourhood | 4 | 27.14 | <0.0001 |
| Replicate | 4 | 0.94 | 0.92 |

Thus, for a given ratio, resistance spreads to a higher frequency when the size of the neighbourhood increases, in a similar manner to what was observed with a lone constitutive response (Figure 1). Intermediate values of the ratio allow the spread of resistance as well as the maintenance of the parasite (Figure 4).

**Figure 4.**
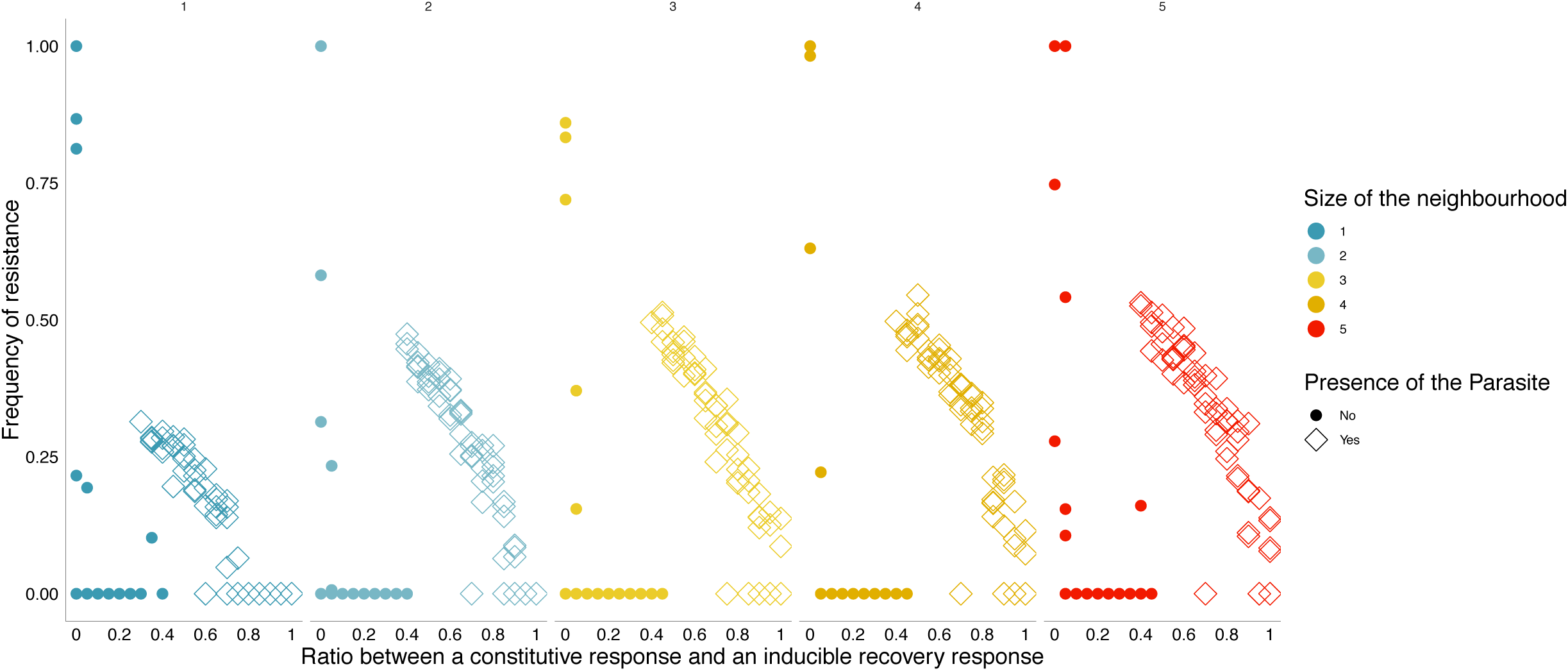
Effect of the ratio between a constitutive and an inducible recovery-boosting resistance on the spread of resistance at different sizes of neighbourhood. The graph presents also the parasite presence at equilibrium. The parameters are the following ones: transmission rate β=0.25, virulence of the pathogen alpha=0.25, mortality μ=0.01, recovery rate γ =0.1, reproduction rate rH=0.02; departure from linearity m1=m2=1.4; maximal cost of the constitutive resistance cMax1=0.1; maximal cost of the inducible defence cMax2=0.1; ratioEff2=1.

Values of r1 close to 1 (corresponding to a higher investment in the constitutive response) permit the spread of resistance only for large neighbourhood sizes without parasite elimination. In other words, when both types of response can occur along a gradient, only the fully transmission-blocking one can become fixed in the population and only a strong investment in it can lead to the eradication of the parasite. If this investment is associated with a fixed cost (due to the constitutive part of the resistance), then resistance is eliminated as well.

#### 3.3.2. Mixing a constitutive and a transmission-blocking response (0 < r1 < 1 and r2=0)

In this scenario, hosts can invest in both a constitutive defence and an inducible response blocking forward transmission (r2=0). As shown in Table 10, both the ratio and its interaction the size of the neighbourhood have an impact on the presence of resistance.

**Table 10.** Logistic analysis of the spread (presence/ absence) of a response with a constitutive component and a transmission-blocking one in different spatial contexts

| Source | d.f. | Log-likelihood $\chi^2$ | P-value |
| --- | --- | --- | --- |
| Size of the neighbourhood | 4 | 7.06 | 0.13 |
| Ratio between the constitutive and the inducible transmission blocking response ($r_1$) | 1 | 22.94 | <0.0001 |
| $r_1$ x Size of the neighbourhood | 4 | 68.23 | <0.0001 |
| Replicate | 4 | 2.38 | 0.67 |

When the value of r1 remains below a certain threshold (i.e. a higher investment in the inducible, transmission-blocking response) resistance can spread for the small-size neighbourhood situations and in a very limited number of situations this can lead to the eradication of the parasite. For a higher investment in the constitutive response (higher values of r1), resistance does not spread except for large neighbourhoods where the parasite is maintained. (S3 Figure).

#### 3.6.3. Mixing all three resistance mechanisms (0 < r1 < 1 and 0 < r2 < 1)

Finally we investigate scenarios in which hosts can invest in all three mechanisms of resistance, i.e. cases where both *r1* and *r2* take intermediate values between 0 and 1. We find that the outcome of the simulations (in terms of presence or absence of resistance at the end of a simulation) is primarily affected by the ratios r1 and r2 of the investment in the different types of the response, the size of the neighbourhood, and its interaction with r1 (Table 11).

**Table 11.** Logistic analysis of the spread (presence/ absence) of a response with a constitutive component, a transmission-blocking and a recovery-boosting response in different spatial contexts.

| Source | d.f. | Log-likelihood $\chi^2$ | p-value |
| --- | --- | --- | --- |
| Size of the neighbourhood | 4 | 63,64 | <0.0001*** |
| Ratio between the constitutive and the inducible transmission response (r1) | 1 | 40,92 | <0.0001*** |
| Ratio between the 2 arms of the inducible responses (r2) | 1 | 17,60 | <0.0001*** |
| r1*r2 | 1 | 2,05 | 0.1519 |
| r1* Size of the neighbourhood | 4 | 145,14 | <0.0001*** |
| r2* Size of the neighbourhood | 4 | 1,38 | 0.8484 |
| r1*r2*size of the neighbourhood | 4 | 3.94 | 0.4144 |
| Replicate | 4 | 0.56 | 0.9676 |

For the different sizes of neighbourhood, 3 different outcomes can occur (Figure 6).

For low values of r1, the fixed cost of resistance is limited and resistance can spread to a high frequency leading to the elimination of the parasite. If there is a minimal investment in the recovery-boosting response, resistance can even fully invade the population and replace the susceptible population. At intermediate values or r1 (approximately 0.1 < r1 < 0.2) resistance is always lost from the population, in association with the fact that parasites are also lost in nearly all simulations (except when the inducible response is almost strictly affecting transmission). For higher values of r1, while resistance can be maintained in the population, we observe that the frequency of resistance is generally lower in small neighbourhoods. Additionally, in these scenarios, higher values of r2 appear to be favoured when r1 is itself higher. In other words, an investment in a recovery-boosting inducible response is favoured when the investment in a constitutive immune response is itself high. In any case, it is interesting to note that, for every neighbourhood size, the highest frequency of resistance was always observed for an intermediate value of r2, suggesting that the combination of both inducible defence mechanisms is most advantageous. Finally, we note that at very high values of r1, resistance is always lost from the population.

## 4. Discussion

Host resistance to pathogens can be based on a variety of mechanisms and investment strategies, and the respective selective pressures associated with these different types of resistance can be correspondingly diverse. In this study, we elucidate how these mechanisms of resistance can be differentially selected, not only based on the way the response to a pathogen is mounted (inducible or constitutive resistance), but also based on the specific trait of the host-pathogen interaction affected by this response.

If we consider specifically the scenarios resulting in fixation of resistance (in the sense of 100% of the population being resistant), we observe that such a fixation never occurs for an allele conferring only constitutive resistance, except when no cost is associated with resistance. This corroborates previous theoretical studies dealing with polymorphism of resistance [9, 10] as well as experimental results with plant species infected by rust fungi [11]. The existence of a negative feedback between the prevalence of resistant hosts and their fitness advantage may indeed limit the evolution of resistance [11, 12].

For the inducible defence, our study shows that fixation can occur not only for a non-costly one but also when the cost remains below a certain threshold. This is valid both for an altruistic and a recovery-boosting resistance. When the resistance is affecting transmission, it can only lead to fixation for small neighbourhood situations for limited values of the cost. When the defence mechanism boosts recovery, fixation can occur at higher costs and for the different sizes of neighbourhood. This is obviously an advantage over a constitutive resistance mechanism. Similarly, one might speculate that transmission-blocking resistance mechanisms, since they do not protect the host from infection, create conditions that are comparatively more detrimental to susceptible competitors, offering an additional benefit. This is reminiscent of comparable effects that have been discussed in the context of the evolution of tolerance [13].

In broad evolutionary terms, the two types of inducible defence considered in this study (increasing recovery and blocking transmission) could be viewed in relative terms as egoist and altruistic strategies respectively. It is beyond the scope of this study and beyond the purpose of this model to draw conclusions about the evolution of altruism from the results presented here, most notably because this model does not keep track of the genetic structure of the population, of relatedness between neighbouring individuals, or of genotypic interactions between hosts and pathogens. Nevertheless, the easy spread of the transmission-blocking response in a small neighbourhood highlights the importance of space in the evolution of altruistic strategies. Such strategies counterselected or selectively neutral in a non-spatial setting, like suicide upon infection [6] can be selected and even become fixed or at least maintained when interactions are limited to their nearest neighbours as shown previously [14] with selection acting through inclusive fitness benefits for the host. Our results also corroborate previous works where altruistic individuals could be maintained under restricted local interactions despite the evolutionary response of egoists [15].

Importantly, the success of strategies based on a mixture of constitutive and inducible responses (e.g. Figure 5) highlights the possible emergence of a resistance polymorphism, despite the presence on a single parasite (as opposed to a community of parasites infecting the host, which is likely to be the norm in natural systems). In addition, our model does not allow any polymorphism in the parasite population, which could favour cycles of co-evolution between hosts and parasites [16, 17]. Moreover, under those conditions where investment is split between these two mechanisms, both resistance and parasitism are maintained in our simulations. Interestingly our work also reveals the possibility that, under certain conditions, a system mixing several resistance alleles can be favoured, and result in both the elimination of the pathogen and the loss of the resistance system itself. Such a system would of course remain open to a later reintroduction of the parasite, as it leaves the host population uninfected and susceptible.

**Figure 5.**
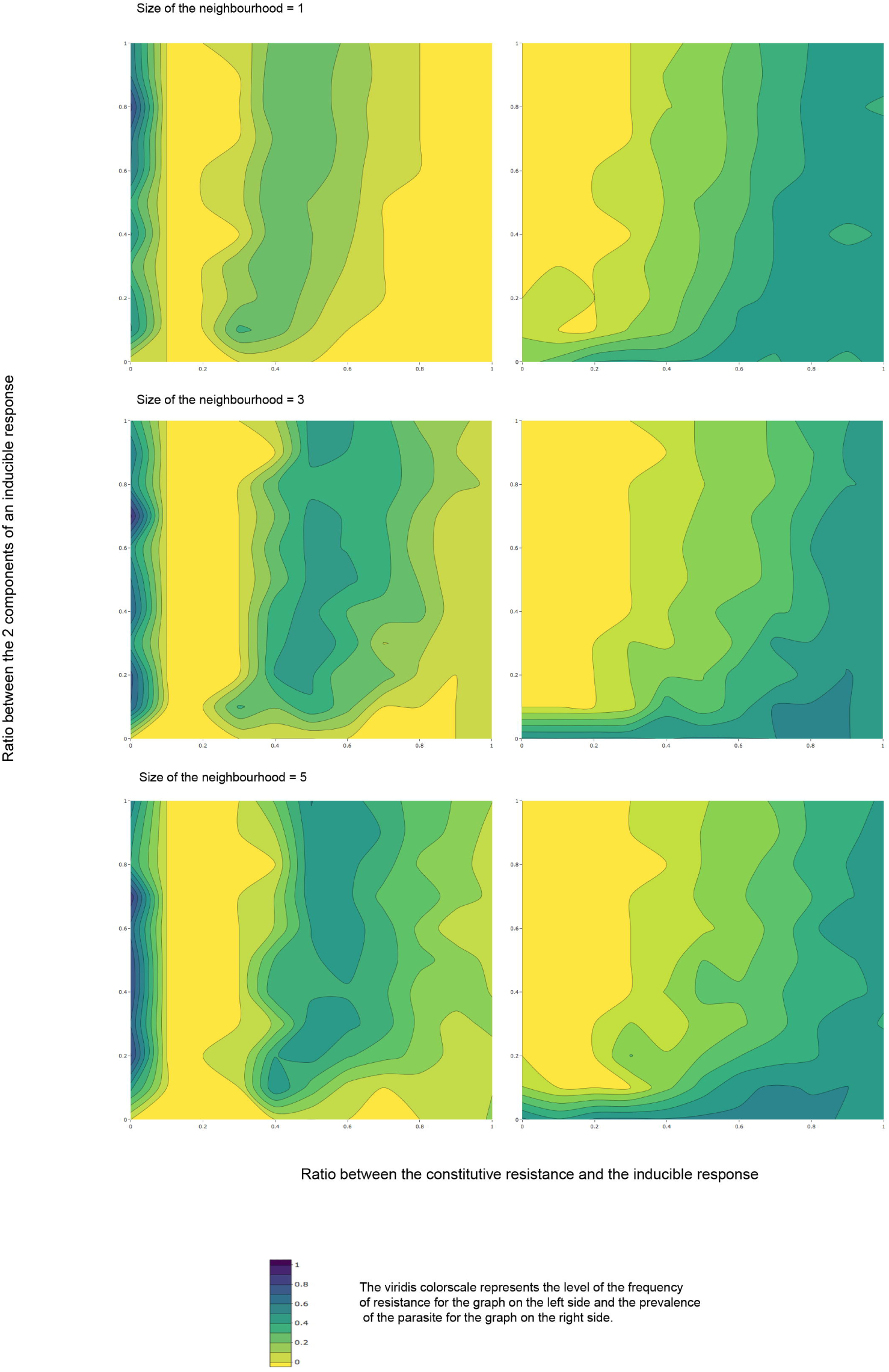
Effect of the ratios between the different types of resistance on the spread of resistance and on the parasite prevalence for different sizes of neighbourhood. Each graph represents the average values of the measured parameters, respectively the frequency of resistance and the prevalence. The parameters are the following ones: transmission rateβ=0.25, virulence of the pathogen alpha=0.25, mortality mu=0.01, recovery rate γ=0.1, reproduction rate, rH=0.02; departure from linearity m1=m2=1.4; maximal cost of the constitutive resistance cMax1=0.1; maximal cost of the inducible defence cMax2=0.1.

As with any model, whether mathematical or not, our assumptions and equations cannot fully capture the complexity of the epidemiological processes. Our results derive from stochastic simulations and as such they represent a complementary approach to the existing literature that is mostly relying on a combination of analytical and simulation results [5, 6, 14]. While many epidemiological models are formulated in continuous-time, our model is formulated in discrete-time. This approach has received less attention but has been used nonetheless in theoretical studies on the population dynamics of infectious diseases [18], on the evolution of immunity [19] or on disease resistance and tolerance in spatially structured populations [7].

For the sake of simplicity, we have made a number of assumptions that inevitably might be considered artificial compared to a number of natural systems. One such assumption is our choice to have all biological processed operate at the same spatial scale (with the same neighbourhood size) in the model. While this offers the advantage of simplicity, it also corresponds to situations already encountered in theoretical system [7] as well as in experimental and field studies with species or population with high or low dispersal. Similarly we have limited the number of parameters in our model with regards to the description of the interaction between hosts and pathogens, and our exploration of the associated parameter spaces. In some cases this was done to focus the analysis, for instance with the parameters associated with the departure from linearity in the cost structure (m1 and m2). We chose to restrict the parameter space to a single value for these parameters since preliminary analyses showed no qualitative differences on the outcome of simulations related to a constitutive response. In other cases we had to make arbitrary choices to limit the scope of our exploration. As an example, we chose to have costs of resistance affect host fecundity (as it was presumably more relevant in the context of local density-dependence within a neighbourhood) but we recognize that the dynamics might differ if the costs of resistance were to affect host survival instead.

Another notable point is that we have not been considering any coevolutionary response of the parasite. This would obviously be an important driver of the dynamics of any epidemiological system but this would require adding complexity to this model. The absence of different types or strains of parasites could also be considered as a default in our model as this would probably favour the selection of resistance polymorphism. There is little doubt that the dynamics of resistance would be affected by the presence of parasites harbouring different life-history traits but this is would add the complexity of interactions between parasites themselves. Similarly, we have not been considering the individual variations in infectiousness, whose role in disease emergence has been established [20]. The presence of superspreaders in particular would likely affect the epidemiology and add complexity to the selection of resistance. Such expansions to the model remain to be done in further studies to better understand the coevolutionary dynamics of host-pathogen systems.

One important caveat specifically associated with our choice of model structure is that the levels of pathogen pressure can vary substantially among our simulations. This could indeed be considered problematic as the infection risk has been shown to favour the constitutive defence mechanism over the inducible one in bacteria [21]. Given our assumptions regarding parasite transmission, the force of infection on an infected host is, on average, higher in larger neighbourhoods. As we believe that it reflects a realistic characteristic of spatially structured populations we chose not to arbitrarily constrain this force of infection, but rather to track prevalence throughout our simulations along with the proportions of sensitive and resistant hosts. Consequently, we did observe some variation in prevalence levels among simulations. This means that the results presented here are driven not only by host-pathogen evolutionary dynamics, but also by host-only evolutionary dynamics in many cases. In other words, these simulations describe the evolution of resistance more at an epidemiological time scale, where wild-type and resistant strategies coexist and compete within a population under a diversity of selective pressures rather than depict the evolution of defence strategies at a true evolutionary time scale.

The epidemiological scale chosen in the current study is nevertheless particularly relevant for many issues regarding host-pathogen interactions and host resistance to disease. The spatial scale of disease transmission is indeed a key factor for the characterization of ongoing epidemics for many diseases e.g. dengue [22-24] or Ebola [25]. Information regarding this spatial scale is often critically important for decision makers regarding the scope of control measures to adopt in the fight against disease transmission [26-28]. Finally, the spatial aspects of resistance strategies and host population dynamics are likely to be crucial to the outcome of strategies based on the use of genetically modified organisms to combat infectious diseases [29, 30].

To conclude, our approach in this study has been to consider the evolution of various defence strategies, alone or in combination, in a spatial setting. As this model was designed to simulate the dynamics of generic host-parasite interactions, the results presented here should not be seen as practical lessons for any particular host-pathogen system. Nevertheless, this study highlights several essential points for a better understanding of the evolutionary dynamics of such systems. First, it calls for the need to consider spatial interactions to understand the maintenance of costly host defence strategies that potentially could not persist in a non-spatial system. Second, as a corollary, it emphasises how essential it is to better understand the maintenance of polymorphism in resistance mechanisms in natural population and the epidemiological dynamics of host-pathogen interactions.

The results of this study could also be informative for more applied purposes. This approach could indeed be useful in defining the desired characteristics of an engineered resistance that could be used in a pest control strategy relying on an added resistance mechanism [31], which would have to cope with the already existing natural immune responses of the host (e.g. genetically modified mosquitoes able to resist malaria). Moreover, considering an engineered resistance based on a mixed-strategy would have several advantages. First, it should limit the evolution of parasites able to avoid the resistance mechanism. Second, it could permit to get rid of the parasite while simultaneously avoiding the maintenance of an engineered resistance in the population. Thus in agricultural systems, this could inform about the best strategies for protective measures of plants through the optimization of the use of one or several cultivar resistant to a pathogen (as discussed for example in [32]).

Overall, it appears clearly that taking into account the spatial structure of host-parasite system is essential to better understand the causes that lead to host heterogeneity in nature [33, 34]. Moreover, and as shown previously [34], the contrasting outcomes between various spatial structures are clearly depending on the cost of resistance but also on the type of response against a pathogen. Concerning the question of cost, the existence of a density-dependant reproduction leads to a lower fitness cost in a spatial setting as shown previously [36]. In that context, our results should permit to better understand the spatial, epidemiological and evolutionary dynamics of infectious diseases, while also highlighting the need for further experimental work to elaborate efficient and sustainable strategies for disease management.

## Acknowledgements

The authors are grateful to Roland Regoes and Samuel Alizon for helpful comments on an earlier version of the manuscript and to King Li for his technical assistance.

## Supporting information

**S1 Figure**. Effect of the fitness cost of resistance and of the size of neighbourhood on the spread of a transmission-blocking resistance against a pathogen and on the parasite presence at equilibrium.

The parameters are the following ones: transmission rate β=0.25, virulence of the pathogen alpha=0.25, mortality μ=0.01, recovery rate γ=0.1, reproduction rate rH=0.02, efficacy of resistance eff2=1, departure from linearity m2=1.4, level of investment in the constitutive resistance ratio1=0, level of investment in the recovery resistance ratio2=0.

**S2 Figure**. Effect of the fitness cost of resistance and of the size of neighbourhood on the spread of a resistance increasing recovery against a pathogen and on the parasite presence at equilibrium.

The parameters are the following ones: transmission rate β=0.25, virulence of the pathogen alpha=0.25, mortality μ=0.01, recovery rate γ=0.1, reproduction rate rH=0.02, efficacy of resistance eff2=1; departure from linearity m2=1.4, level of investment in the constitutive resistance ratio1=0; level of investment in the recovery-boosting resistance ratio2=0.5.

**S3 Figure**. Effect of the ratio between a constitutive and an inducible transmission-blocking resistance on the spread of resistance at different sizes of neighbourhood. The graph presents also the parasite presence at equilibrium.

The parameters are the following ones: transmission rate β=0.25, virulence of the pathogen alpha=0.25, mortality mu=0.01, recovery rate μ=0.1, reproduction rate, rH=0.02; departure from linearity m1=m2=1.4; maximal cost of the constitutive resistance cMax1=0.1; maximal cost of the inducible defence cMax2=0.1; ratioEff2=0.

